# Coverage, Opportunity and Challenges of expanded Program on Immunization among 12-23 Months old Children in Woldia Town, Northeast, Ethiopia, 2018: Mixed cross sectional study

**DOI:** 10.1101/479311

**Authors:** Ayele Mamo Abebe, Mesfin Wudu Kassaw, Alemu Birara

## Abstract

**Objective:** The purpose of this is to assess coverage, opportunity and challenges of EPI among children age 12-23 month in Woldia town, Amhara region, Ethiopia.

**Result:** A total of 389 mothers/caretakers were interviewed. Based on vaccination card and mothers/caretakers’ recall, 385 (99%) of the children took at least a single dose of vaccine. About 343 (87.7%) children were fully immunized. Dropout rate was 9% for BCG to measles. Qualitative study revealed that workload, shortage of vaccine and non-compliance of mother/care taker for next schedule date was the major challenge faced by health professionals. In this study, vaccination coverage was low compared to the Millennium Development Goals target. Thus the town health office and concerned stakeholders need to work more to improve performance of the expanded program on immunization in this area.

## Introduction

Expanded Program on Immunization (EPI) began by World Health Organization (WHO) in 1974[1,2]. About 107 million babies (83%) global had gotten at least 3 doses of DTP vaccine; but, nearly 22.4 million miscarried to acquire 3 dosages, exposing huge numbers of children susceptible to vaccine-preventable diseases and death [3,4]. One out of five infants worldwide does not receive 3 life-saving doses of the diphtheria, tetanus and pertussis vaccine [5].

Below sub-Saharan African, 4.4 million children were die yearly due to transmittable diseases that could be avoidable by immunization [6,7]. It is associated to poor immunization Coverage, challenges and setup were not fully equipped in sub-Saharan Africa countries [8].

EPI was began in 1980 in Ethiopia with the purpose of reducing decease and disease of children and mothers from vaccine avoidable diseases [9]. To achieve maximal protection against vaccine-preventable diseases, a child should receive all vaccines[10, 11,12, 13]. It has been also recognized that vaccine preventable diseases are responsible for 16% of under-five mortality in Ethiopia [14]. Even if Ethiopian government did different efforts, but the coverage rates was stayed highly small for several centuries [16,17].

AS explained by EDHS 2016, two from five children aged 12–23 months (39%) had gotten totally fundamental immunizations at some time, also 22% have gained vaccine by the suitable period time. There was increment of fully vaccinated children aged 12–23 months percentage from 24% to 39% [18]. Based on the EDHS 2016 report, there is a wide difference among regions concerning full immunization coverage ranging from 89% in Addis Ababa to 15 % in Afar region due to different challenges and opportunities as explained in other studies [16, 17, 18, 19& 20].

But in Woldia town there was no previous study that shows the coverage, dropout rate of EPI and asses’ different challenge and opportunity of EPI. Therefore the aim of this study is to determine the level of immunization coverage, dropout rate and to explore challenges and opportunity of EPI among all pair of mother to children aged 12-23 months in Woldia town.

## MAIN TEXT

### Study Area and Period

A community based cross sectional study using mixed method was conducted in Woldia town from April 26-May 11, 2018. Woldia is the capital of the north Wollo zone which is located at about 520 km away from Addis Ababa capital city of Ethiopia. According to 2007 Ethiopia national census the total number of children age 12-23 months is 2250(38). Woldia has 1 general hospital, 2 health center, and four health post. The Source of population was all children aged 12 to 23 months with their mothers/caretakers living in Woldia town.

### Sample size calculation

The sample size was calculated using the single population proportion formula based on the following assumption. Using p = 76% **(22)**, design effect (DE) =1.5 and CI=95%

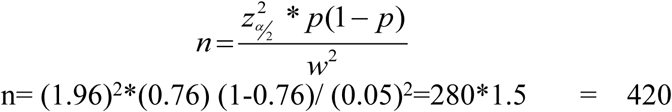

Our total population is< 10,000 so; we use the following correction formula

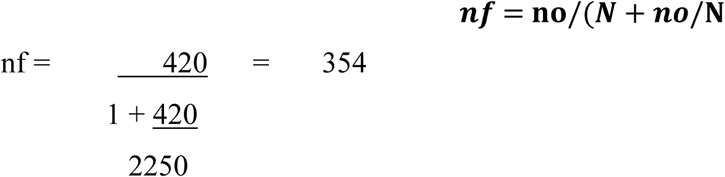

By adding 10% non-response rate we get a final sample size is 389

### Sampling technique

Multistage sampling was used to get the sample. Four kebeles were selected from 10 kebeles by simple random technique. The total sample size was allocated proportionally to each kebeles depending on the total number of children. The first house hold had been selected randomly from selected kebeles. The sample for in-depth interview was selected by non-probability purposive sampling method. A total of 11 in-depth interviews were conducted among health professionals and HEW who works under EPI programs in Woldia town (**Fig 1**).

**Figure 1.**
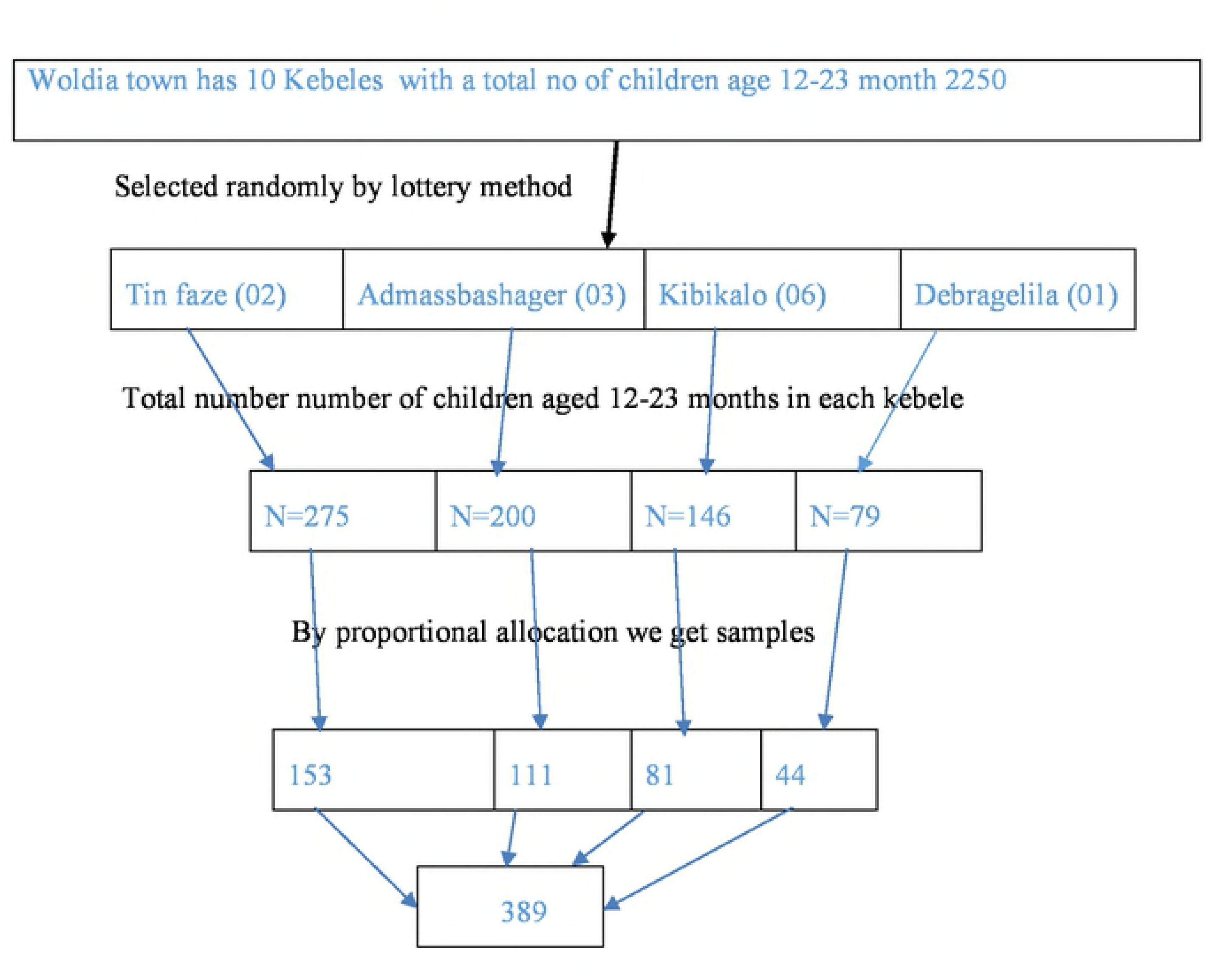
schematic presentation of sampling procedure

### Operational definition

#### Fully vaccinated

children are considered as fully vaccinated when they have received all ten vaccines (19, 20).

#### Vaccinated

children are considered as vaccinated when they took at least one dose of the vaccines (19, 20).

#### Dropout rate

It is the rate difference between the first and the last dose (19, 20).

#### Coverage by Card Only

It means coverage made with numerator based on only recorded dose.

#### Coverage by Card plus History

It means coverage done with numerator based on card and mother’s report (19, 20).

### Data collection tool and Procedure

The questionnaire was developed from previous studies (16, 20, and 17). In-depth interview schedule will be used for collect qualitative data. The interviews were documented appropriately by participant words and the participants speaking was recorded by tape and transcribed to the English word.

### Data quality and control measure

The questioner was prepared by English language and translated to Amharic language and again translated to English by professional translators. Pretest was done in 5% of sample size at similar setting near in the study setting (defergie Keble) and necessary modification was done. Training was given for two day about data collection. Completeness of data were checked at the end of each day of data collection by principal investigators and supervisors.

### Data processing and Analysis

Data were coded, entered and analyzed using SPSS version 20. Result was presented by table, frequency and graphical techniques were used. Recorded or paper documented file from the qualitative data were transcribed into Microsoft word 2010. Similar topics were grouped together, and those with common features were clustered together until the final themes and sub-themes emerged.

## RESULT

### Sociodemographic Characteristics of the participants

A total of 389 mothers/caretakers were interviewed with a response rate of 100%. The majority 195 (50.1%) were from age of 25 and 31. The immediate caregivers of the children were mothers (97.9%).About, 84.8% of the caregivers were currently married. With regard to religion, 324 (83.3%) were orthodox. About 371(95.4%) respondents were Amhara in ethnic group. Regarding educational status, 54.7% participants have secondary and above. By occupation 226 (58.1%) were housewives. Relating to income, 258 (66.3%) participants had monthly income greater than 500 birr (**Table.1**).

#### Family size and characteristics of the child

Among the respondents 171 (44%) have one child, and 161(41.4%) have 2 or 3 children. The most family size of the study participants was 4 ranging from 2 to 9 (70.4%). The mean age of the children was 17.38 months and 218 (56%) were of the male gender (**Table 2**).

#### Immunization Coverage of Children less than two years

Three hundred forty one (87.7%) of children completed all the suggested doses. Using card only, 98 (25.2%) children had vaccination card. From 98 vaccinated by card only, 98.9% received OPV1. Measles and Penta 3 vaccines were taken by 72.4% and 86.7%, respectively. According to card plus mother recall, 385 (99%) of the children had taken as a minimum a single dose of vaccine. However, 341 (87.7%) were claimed fully immunized (**Additional table 1**).

### Dropout Rate for Vaccines

According to the findings from the field survey in the number of children who defaulted on the vaccines from DPT1 and OPV1 to Measles. Forty two children (10.8%) defaulted for Measles getting from both recall and card. The penta1-measle dropout rate for children was 8.3% and penta1-penta3 and pcv1-pcv3 dropout rate was 2.4% and 1.6% respectively .the overall dropout rate (from BCG-Measles) was 9%.

### Reasons for Defaulting from Vaccination service

In this study, 2.3 % of respondents reported that the reason for not completing child vaccination was lack of awareness about vaccination and same 2.3% not knowing come back for second and third vaccination, 2.1% fear of side effect, 1.8% vaccination inconvenient time and no vaccination at health facility for each(**Additional table 2**).

### In-depth interview findings

The purpose of this in-depth interview was to explore challenge and opportunity of EPI and also how those challenges relieved. Themes and sub-themes were used that reflect idea of participant.

#### Theme 1: challenges faced by HW and HEW

HEW and health professionals highlighted that they faced challenges when implementing EPI those are shortages of vaccines, workload, and Non-compliance of mothers with scheduled return dates.

##### *Subtheme 1.1*: Shortage of vaccines

Health workers had explained as they have sometimes experienced a scarcities of vaccines.

##### Subtheme 1.2: Workload

Participants explained that workload decreases the accuracy of records of performance on vaccination and reduces chances of counseling clients on the importance of vaccines.

##### *Subtheme1.3:* Non-compliance of mothers with scheduled return dates

According this study, most mothers have not knowledge which diseases are prohibited by vaccines, or needed doses of each vaccine is not known by them.

#### Theme 2: Possible solutions to relieve faced challenges

The possible solutions were prepared enough work area, sufficient amount of vaccine, having enough staff, more health education about when to return, proper use of card and what vaccine at what age of child give and what type of disease prevented by that specific vaccine.

#### Theme 3: opportunity that increases EPI Coverage

##### Subtheme 3.1: Health professional’s awareness regarding internal referral system

As confirmed by most study subjects who said: “most health professionals are aware of the internal referral system. Most health professionals also advise mothers who come for maternity services to vaccinate their child”.

##### Subtheme3.2: information accessibility

As confirmed by most participants said; “our community gets information regarding immunization easily from media, HEW and other sources since it is urban community.

##### Subtheme 3.3: outreach service delivery

HEW said that all kebeles have health extension worker so, the HEW round and searching unvaccinated and partially vaccinated children’s if get give a vaccination for child and appoint for next schedule mothers go to near health facility and also give education regarding what health problem is occurred if the child is completed vaccination.

## DISCUSSION

This study was conducted to assess coverage, opportunity and challenges of EPI among children in Woldia town, Amhara regional, Ethiopia. In this study,each vaccine‘s Coverage had be 96.9% for BCG, 98.2% for OPV1, 97.4% for OPV2,94.6% for OPV3, 96.2% for pentavalent1, 96.9% for pentavalent2, 93.9% for pentavalent3, 95.9% for pneumococcal conjugated vaccine (PCV1), 97.4% for PCV2, 94.3% for PCV3, 96.4 for Rota1, 94.8 for Rota2 and 88.2% for vaccine of measles. The finding of this study was higher than the previous study done in different areas [20, 21, and 22]. This may be due to improving vaccination’s access and awareness of a community from time to time. But this study was lower than study done in Debre Markos town (91.7%) [23]. This may be due to insufficient vaccine available.

The overall dropout rate for this study was 9%. Dropout rate from pentavalent1 to measles (8.3%) was greater than dropout rate from pentavalent (2.4%).However, this finding was lower than other studies such as EDHS 2011 and a study done in Yirgalem town [15, 20]. On the other side, this study has a higher dropout rate from study done in Debre Markos town (5%) [23]. This may due to workload, insufficient of vaccine.

In this study, the percentage of fully vaccinated children was 87.7% which showed that there had be significant amount of non-attending children. Participants mentioned different causes for not finishing vaccination to children. Those causes were lack of awareness about immunization, not knowing coming back for second and third vaccination, fear of side effect, inconvenient vaccination time and insufficient vaccine. This finding was supported by other similar studies. [20].

In qualitative study, workload due to staff shortage and inadequate workspace, shortage of vaccine and non-compliance of mother for next scheduled date were the major challenges faced by health professional and health extension worker. This was aligned with a study done in Arebegona district Southern Ethiopia [24].

## Conclusions

This study was lower the nationwide MDG target (90.0%). Reasons for incompletion are lack of awareness about immunization, not knowing coming back for other schedules and fear of side effect. Reason for not vaccinating their child, most respondents replied that lack of awareness on importance of vaccination, fear of side effect and sickness of child as a reason.

Work load, non-compliance of mother and shortage of vaccine was the major challenges faced by health professionals and health extension workers.

### Limitation of study

Since the study was cross sectional it did not shown the cause-effect relationship.

## Abbreviations

BCG: Bacillus Calmette-Gurin
DHS: Demographic health survey
DPT: Diphtheria, Pertussis, Tetanus
EC: Ethiopian calendar
EDHS: Ethiopian Demographic and Health survey
EPI: Expanded Program on Immunization
GVAP: Global vaccine action plan
HEB: Hepatitis B
MCV: Measles containing Vaccine
MDG: Millennium development goal
MOH: Ministry of health
OPV: Oral Polio Vaccine
PCV: Pneumococcal Conjugate Vaccine
SNNPR: Southern nation nationality people region
SPSS: Statically package for social science
UNICEF: United Nations International Children’s Emergency Fund
VPD: Vaccine preventable disease and WHO: World Health Organization

## Declarations

### Ethics approval and consent to participate

Ethical clearance was obtained from Woldia University, ethical review board and permission was given from Woldia town health administrative office and administrative of selected kebeles. The research proposal was evaluated and approved by the Research Ethics Review Committee [HRERC 0610/2018] of College of Health Sciences, Woldia University and ethical clearance was obtained. Official cooperation and permission was obtained from Woldia town health administrative office and selected kebeles. Moreover, prior to commencing the study, a written informed consent was obtained from each respondent before data collection. Written consent was found from a respondent. Confidentiality was maintained by omitting their name and personal identification of participant was no compelled to the study. This means Confidentiality was not shared for others.

### Consent for publication

Not applicable.

### Availability of data and materials

Data supporting the conclusions of this article are available by request to Ayele Mamo. The relevant raw data will be made available to researchers wishing to use them for non-commercial purposes. The Supplementary data will be put below the acknowledgement.

### Authors’ contributions

All authors wrote the research, developed the questionnaire, analyzed the data and wrote the paper and interpreting of the findings as well as participating on the preparation of the manuscript. MW and AB supervised the data collection, contributed to the interpretation of the findings. AM and MW trained data collectors, participated on the preparation of the manuscript. All authors read and approved the final manuscript.

## Acknowledgement

The authors would like to acknowledge Woldia university financial support and Woldia University for providing sponsoring ship. Woldia town health Administrative office and all study participants in selected Kebeles are acknowledged for their cooperation during sample collection.

---

Figures and tables legend for Coverage, Opportunity and Challenges of expanded Program on Immunization among 12-23 Months old Children in Woldia Town, Northeast, Ethiopia, 2018.

| S.no | Description of tables and figures |
| --- | --- |
| 1 | Table 1: Socio demographic characteristics of caregiver in Woldia Town, North wollo zone, North East Ethiopia, May, 2018(n=389). |
| 2 | Table2: Family size and characteristic of child, in Woldia Town, North east Ethiopia, May, 2018(n=389). |
| 3 | Figure1: schematic presentation of sampling procedure |

Additional tables legend for Coverage, Opportunity and Challenges of expanded Program on Immunization among 12-23 Months old Children in Woldia Town, Northeast, Ethiopia, 2018.

| S.no | Description of additional tables |
| --- | --- |
| 1 | Additional Table 1: Vaccination Coverage of children aged 12–23 months based on child vaccination card and mothers recall, in Woldia town, North Wollo Zone, Northeast Ethiopia, 2018(n=389). |
| 2 | Additional table2: Reasons given by caregivers for not completing children Vaccination, Woldia town, Northeast Ethiopia, May, 2018(n=389) |

